# Single-cell profiling reveals disruption of cell-cycle homeostasis following JDP2 depletion in colorectal epithelial cells

**DOI:** 10.64898/2026.09.14.751340

**Authors:** Violet Newhart, Angela Gao, Ana Edens, Madison Flori, Paloma Bravo, Ashfaqul Alam

## Abstract

Intestinal epithelial cells (IECs) undergo rapid and continuous renewal to maintain gut epithelial homeostasis and barrier integrity at the interface with luminal microbiota, dietary antigens, and enteric pathogens. IEC proliferation is tightly regulated by a complex machinery of cell cycle regulators and effectors, and genetic or transcriptional dysregulation of cell-cycle regulatory pathways is a hallmark of colorectal cancer development and progression. Previously, JDP2 (Jun dimerization protein 2) has been shown to be involved in cell cycle control and is associated with an array of cancers. JDP2 functions as a transcription factor or epigenetic regulator depending on cellular context, disease, and cancer types. However, JDP2’s role in CRC, cell cycle homeostasis of colon cancer cells, and their fate remains poorly defined. In this study, we found that patients with higher JDP2 expression have significantly worse survival than patients with lower JDP2 expression. Our functional assays showed that JDP2 depletion increased EdU incorporation during S phase and PHH3 abundance, which is consistent with increased proliferative and mitotic activity. Our single-cell RNA sequencing and transcriptomic profiling further revealed reprogramming of cell-cycle-associated transcriptional states following JDP2 depletion. Differential-expression and pathway analysis identified changes in mitotic and cell-cycle regulatory genes, including programs involving cyclins, cyclin-dependent kinases, chromosome segregation, and mitotic progression. We also found that lower residual JDP2 expression was associated with escalated G2/M representation and a change toward later states along a G1→S→G2/M-associated transcriptional trajectory. Furthermore, low-JDP2 cells showed significantly greater late-state occupancy than High-JDP2 cells (33.6% versus 24.0%; paired P=0.048). Among the JDP2-siRNA-treated cells, analysis of residual JDP2-expressing cells also revealed heterogeneous associations of JDP2 with cell-cycle, survival, apoptotic, and stress-response programs, consistent with JDP2’s context-dependent transcriptional functions. Together, these data identify JDP2 as an important component of intestinal epithelial cell-cycle homeostasis and transcriptional landscape and suggest that JDP2 depletion perturbs the balance of proliferative cell states.

## Introduction

The intestinal epithelium is a highly dynamic tissue that undergoes continual renewal, proliferation, and differentiation of the intestinal epithelial cells (IEC) to maintain gastrointestinal homeostasis. This rapid turnover is a tightly regulated progression of IEC through the cell cycle, balanced by differentiation and epithelial cell shedding. Cell-cycle progression is carefully orchestrated by networks of cyclins, cyclin-dependent kinases (CDKs), CDK inhibitors, transcriptional regulators (TFs), and checkpoint mechanisms. These interconnected regulators coordinate DNA replication, chromosome segregation, and mitosis. Genetic, epigenetic, or transcriptional disruption of cell-cycle homeostasis may lead to uncontrolled proliferation of IECs, a hallmark of colorectal cancer (CRC)^1–4^. Since 2022, colorectal cancer (CRC) has been classified by the WHO International Agency for Research on Cancer as having the 3rd highest incidence rate of any cancer worldwide and the 2nd highest mortality rate^1,2^. The number of CRC cases has been growing since 2020, with 2020 recording 1,931,590 cases globally and estimates of 2040 reaching 3,154,674 cases^5^. In the United States, new CRC cases are projected to increase by 0.21 million by 2040. In the US alone, around 24.3 billion USD was spent on CRC care and treatment^6^. Hence, determining the molecular mechanisms that modulate IEC proliferation and the cell cycle is critical for understanding CRC pathogenesis and identifying molecular determinants of tumor development and progression.

Over the years, a body of work has shown that CRC develops through the progressive accumulation of genetic and epigenetic alterations that may disrupt signaling networks, which regulate epithelial proliferation, survival, differentiation, and genome stability^4^. CRC may arise through multiple, partially overlapping routes of tumorigenesis. These include the conventional adenoma–carcinoma sequence and the serrated neoplasia pathway, accompanied by molecular alterations such as chromosomal instability (CIN), microsatellite instability (MSI), and aberrant DNA methylation. In colorectal cancer, APC/WNT, KRAS/MAPK, TP53, PI3K, TGF-β/SMAD, and DNA mismatch repair are the most frequently altered pathways^4–8^. Although they have distinct molecular functions, these pathways converge into the core networks that regulate gene expression and cell-cycle progression^7,8^. Ultimately, the combined effects of these alterations influence whether colorectal epithelial cells continue dividing, pause, or exit the cell cycle altogether. As for an example, loss of APC causes constitutive WNT/β-catenin signaling and aberrant expression of proliferative programs. In addition, alterations in KRAS and TP53 further disrupt growth signaling, checkpoint control, and cellular stress responses. Thus, CRC reflects progressive disruption of the molecular mechanisms that normally coordinate cell-cycle transitions and epithelial cell states. Identifying transcriptional regulators that control these states is therefore important for understanding colorectal tumor biology.

JDP2 (Jun dimerization protein 2) is a bZIP transcriptional regulator that interacts with AP-1 family proteins, including JUN and ATF, and modulates AP-1-dependent gene regulation. Importantly, depending on its binding partners, cellular context, and promoter, JDP2 can act as both a transcriptional repressor and activator. Furthermore, JDP2 is involved in chromatin organization and histone acetylation^7–10^. Functionally, JDP2 participates in cell-cycle control, differentiation, apoptosis/survival, senescence, and oxidative-stress responses, including interactions with pathways involving p53 and NRF2^12–19^. Hence, JDP2 can have cellular state-specific and tumor type-dependent tumor-suppressive or tumor-promoting functions^7–10^. It is not clear how JDP2 regulates colorectal cancer development and progression, especially in colorectal cancer cells^11–14^.

A major feature of CRC is the considerable cellular and transcriptional heterogeneity. Indeed, this heterogeneity exists not only between patients and tumors but also among individual cells within the same tumor tissue. In spite of cells sharing the same major genetic alterations, they can still exhibit distinct transcriptional states, as evident from differences in proliferation, differentiation, or stress responses. Importantly, this diversity in cell transcriptional state comes from a mix of genetic and epigenetic changes, cell plasticity, and interactions with the microenvironment. As a result, the levels of certain regulatory genes can be very different from one cancer cell to another in the same tumor. Bulk RNA sequencing averages results across many cells, which can obscure these individual-cell-level gene expression differences. In contrast, scRNA sequencing examines each cell separately, which enables transcriptional heterogeneity to be resolved at cellular resolution and links specific patterns to broader cell behaviors^37–40^. By examining such within-population variation, we can find connections between gene regulators and cell functions that are missed when looking at average data. Given the cell-state-specific transcriptional and cell-cycle functions of JDP2^20–29^, we therefore posited that heterogeneity in JDP2 expression among colorectal cancer cells may be associated with distinct cellular states. Examining JDP2 at single-cell resolution may reveal relationships between its expression and cell-cycle or other transcriptional states that are masked by population-averaged measurements^30^. One approach is to examine heterogeneity or variation in residual JDP2 expression within the same siRNA-treated cell population, allowing JDP2-associated heterogeneity to be studied without requiring a separate control for this within-population comparison^30–36^.

In this study, we determined the survival probability of CRC patients with high versus low JDP2 expression; the effect of spermine on relative JDP2 expression in Caco-2 cells; and that decreased JDP2 levels increased the proliferation marker PHH3 and the proliferation of colorectal cancer cells. We found that siRNA-mediated knockdown of JDP2 increases PHH3 labeling in CRC cells. Furthermore, depletion of JDP2 increased overall EdU incorporation during the S phase and increased phosphorylation of PHH3. Our scRNA sequencing analysis further demonstrated redistribution of cell-cycle-associated transcriptional states following JDP2 depletion by siRNA. Further differential-expression analysis identified changes in mitotic and cell-cycle regulatory genes, including programs involving cyclins, cyclin-dependent kinases, chromosome segregation, and mitotic progression. Among JDP2-siRNA-treated cells, residual JDP2 expression also revealed heterogeneous associations with cell-cycle, survival, apoptotic, and stress-response programs, consistent with JDP2’s tumor type and cell state-dependent transcriptional functions^20–29^. These findings elucidate a pivotal role of JDP2 in intestinal epithelial cell-cycle homeostasis and suggest that JDP2 depletion disrupts the balance of proliferative cell states.

## Methods

### Caco-2 cell culture and seeding for siRNA transfection

Caco-2 cells cultured in standard MEM media with FBS, PenStrep, Glutamax, and MEM NEAA were trypsinized for 5 minutes and diluted with media to seed in either 24-well plates, 96-well plates, or 8-well chambers slides. Cell seeding numbers were calculated accordingly, and cells were transfected within a range of 60-80% confluence.

### JDP2 siRNA transfection

Standard MEM media and opti-MEM were warmed to 37C in a water bath. The prepared standard MEM media was replaced in all wells. Lipofectamine RNAiMAX Transfection Reagent, JDP2 siRNA, and negative select control were diluted in Opti-MEM. Diluted lipofectamine was added in a 1:1 ratio to the JDP2 siRNA and control siRNA and incubated for five minutes at room temperature. The Lipofectamine-siRNA complex was added to each well and incubated for three days at 37C.

### Protein extraction

After 72 hours post transfection, cells were lysed with RIPA buffer and a tablet of protease and phosphatase inhibitors. Using a 1 mL pipette tip, cells were scraped off the bottom of the 24-well plate and vortexed for 10 seconds. The cells were then incubated for 30 minutes at 4C and subsequently centrifuged for 20 minutes at 12,000 RPM and 4C. The supernatant was removed, and resulting protein was utilized in western blot experiments.

### Immuno blotting

Protein concentrations were measured using a microBCA Protein Assay Kit^42^. Samples were diluted with SDS loading buffer, vortexed, spun down, and incubated at 95C for 5 minutes. Samples were centrifuged for 5 mins at 12000g. 15 *μ*L of protein samples and 7 uL of Westen C Standard protein ladder were added to a 20uL well gel. The gel was run for 5 minutes at 50V and then for 35 minutes at 150V. The transfer step was completed utilizing a Trans-Blot Turbo Transfer System on the mini TGX protocol. The membrane was trimmed and blocked with 1xTBST 5% milk for 1 hour at room temperature. The primary antibody solution was prepared with 1xTBST, 5% BSA, and the membrane was incubated in the primary antibody overnight at 4C. The secondary antibody solution was prepared with 5mL of 1xTBST, 5% milk, 1 µL of StrepTactin-HRP, and goat anti-rabbit secondary antibody. The membrane was incubated in 5mL of secondary antibody dilution for 1 hour at room temperature. Clarity solution was added to the membrane for 5 minutes at room temperature, and the blot was imaged. Band intensity was measured utilizing image j software and statistical analysis was performed to determine the effect of JDP2 siRNA knockdown on JDP2 and PHH3 levels.

### EdU assay and analysis

Cells were seeded in 8-well chamber slides and transfected with 12pmol and 6pmol of control siRNA and JDP2 siRNA. After three days of growth, the EdU assay was performed utilizing the Click-iT EdU Cell Proliferation Kit^41^. EdU labeling solution was added to wells to form a 10uM final EdU solution and incubated at 37C for two hours. Cells were fixed with 4% paraformaldehyde in PBS. Cells were then permeabilized with 0.5% Triton X-100 in PBS for 20 minutes. Next, the Click-iT reaction cocktail was prepared, and the cells were incubated with the cocktail for 30 minutes at room temperature. Finally, cells were stained with a 1 mg/mL DAPI solution for 5 minutes. The slide was coversliped and mounted with prolong gold antifade. After curing at room temperature overnight, the slide was stored at 4C until imaging.

Images were taken utilizing the confocal Nikon A1R microscope at the University of Kentucky Imaging Core. The 403.1 nm and 561.4 nm lasers were used to image the DAPI and EdU channels respectively. Four images were taken per well in different regions of each well. Then, the number of DAPI nuclei and EDU+ cells were manually counted per image. Statistical analysis was performed to determine the effect of JDP2 siRNA knockdown on EdU-associated proliferation.

### RT qPCR

RNA was extracted from Caco-2 cells utilizing the Qiagen RNeasy Mini Kit with an added DNA digestion step. iScript Reverse Transcription Supermix (iScript gDNA Clear cDNA Synthesis Kit, BioRad) was used to prepare the cDNA. qPCR was carried out using SsoAdvanced Universal SYBR Green Supermix (Bio-Rad), gene-specific primers, nuclease-free water, and cDNA. The thermocycler settings were set according to the iTaq kit instructions. Results were analyzed using the ΔΔCt method^41–42^.

### Spermine treatment and transcription-factor screening

To conduct a limited screen for metabolite-responsive transcription factors, we treated Caco-2 cells with spermine for 24 h, followed by RNA isolation and an initial qPCR-based screen of transcription factors using a 96-well plate format (BioRad PrimePCR)^41–42^. We selected transcription factors that showed changes in expression during the initial screener evaluation. Among these candidates, we next examined JDP2 in greater detail by treating Caco-2 cells with increasing concentrations of spermine (0, 10 μM, 100 μM, and 1 mM) for 24 h. Finally, we measured JDP2 expression by qRT-PCR and normalized it to the corresponding housekeeping gene.

### Immunofluorescence (IF) Staining

Caco-2 cells in 8-well chamber slides were fixed with 4% paraformaldehyde in PBS for 20 minutes and blocked with 5% BSA and 0.3% Triton X-100 for one hour. Cells were then incubated with primary antibody dilution buffer overnight consisting of 1% BSA, 0.3% Triton x-100, and rabbit primary antibody. The secondary antibody dilution buffer was prepared with 1% BSA, 0.3% Triton x-100, and Anti-rabbit IgG Fab 2 Alexa Fluor 488 secondary antibody. The cells were incubated with the secondary antibody solution for two hours. Cells were then stained with a 1 mg/mL DAPI solution for 5 minutes. The slide was coversliped and mounted with prolong gold antifade. After curing at room temperature overnight, the slide was stored at 4C until imaging^41–42^.

### Single-cell library preparation and sequencing analysis

Libraries were generated for 3 samples using Chromium GEM-X Flex Fixed RNA Profiling - Human, v2 chemistry with the Chromium Human Transcriptome Probe Set v2, GRCh38-2024-A. We processed all samples in a pooled Flex workflow and demultiplexed them using sample-specific probe barcodes.

We found that the integrated feature-barcode matrix contained 18,132 genes and 90,000 cells. Barcode-level metadata from the 10x graph-based clustering export and condition assignment file matched the matrix exactly, with no duplicated barcodes. Cell Ranger Multi-sample identified 28,337, 32,670, and 30,180 cells in 3 samples. We deposited the single-cell RNA-sequencing data generated in this study in the NCBI GEO, and raw sequencing data are available through the associated SRA.

### Seurat preprocessing, dimensional reduction, and clustering

We performed analyses in R 4.3 using Seurat 5.3 ^37–40^. RNA counts were normalized with NormalizeData (scale factor 10,000), 2,000 variable features were selected, and PCA was performed^30–36^. We also used the first 18 principal components for neighbor graph construction, UMAP, and clustering. Finally, we found that clustering at resolution 1.1 generated 15 transcriptional states (clusters 0–14).

### Cluster markers and quality assessment

Using Seurat FindAllMarkers with min.pct = 0.10, a log2 fold-change threshold of 0.25, and Wilcoxon testing enabled us to identify positive markers for all 15 clusters^37–40^. Then, we examined the cluster-level nCount_RNA and nFeature_RNA distributions.

### Cell-cycle scoring

We also inferred cell-cycle state using Seurat S- and G2/M-phase gene sets; 42/43 S-phase genes and found that all 54 G2/M genes were represented. The current scored object contained G1, G2/M, and S-classified cells. We summarized phase distributions at the condition-, cluster-, and replicate-level.

### Condition-level differential expression

Seurat FindMarkers with the Wilcoxon test was used with logfc.threshold=0 and min.pct=0. We ranked genes by log2 fold change and BH-adjusted p-values. We performed replicate-aware pseudobulk analysis by aggregating raw counts by biological replicate and JDP2 Expression state, then analyzed them with DESeq2 using the paired design ∼ Sample_ID + JDP2_group.

### Pathway enrichment and sample-level pathway visualization

We analyzed replicate aware differential expression lists with ranked GSEA using fgsea against human MSigDB Hallmark and Reactome gene sets. Additionally, we conducted cutoff-free ranking of GSEA with fgsea against human Reactome and MSigDB Hallmark gene sets.

We performed paired pseudobulk comparisons of Low-versus High-JDP2 cells, with analyses performed both across the JDP2-siRNA population, within G2/M cells and clusters. Individual cells can exhibit heterogeneous transcriptional responses within a common genetic perturbation background, as demonstrated in single-cell perturbation studies^20–29,37–40^. Hence, we investigated whether residual JDP2 expression was associated with distinct cellular states among JDP2-siRNA-treated cells. We therefore restricted analyses to three independent JDP2-siRNA biological replicates (S1–S3). Also, we operationally classified cells as Low-JDP2 (log-normalized JDP2 expression ≤1) or High-JDP2 (>1). Next, we applied these categories to interrogate within-treatment transcriptional heterogeneity and did not interpret them as quantitative measures of siRNA uptake or knockdown efficiency. We also utilized cell-cycle, mitotic, checkpoint, apoptosis/stress, and survival programs. We evaluated them using prespecified gene sets. Finally, along the Slingshot cell-cycle-associated trajectory, we summarized standardized module scores across pseudotime separately for Low- and High-JDP2 cells and across the three biological replicates. This was followed by our examination of leading-edge detection frequencies to identify sparse-gene artifacts. We quantified metabolic pathway activity with scMetabolism KEGG gene sets and AUCell in a balanced 10,000-cell subset. We averaged AUCell scores by biological sample.

### Statistical analysis

Functional experiments were performed in at least three independent experiments unless otherwise indicated. Data are presented as mean ± SEM. Comparisons between two groups were generally performed using Student’s t-test, whereas comparisons involving multiple groups were analyzed by analysis of variance (ANOVA), as appropriate. Statistical significance was defined as P < 0.05, with significance levels indicated in the figures or figure legends where applicable (e.g., P < 0.05 and P < 0.01). Statistical analyses for the single-cell RNA-sequencing studies were performed as described in the corresponding Methods sections. Differential-expression analyses incorporated multiple-testing correction using the Benjamini–Hochberg method where appropriate, and replicate-aware pseudobulk comparisons were used to account for biological replication. For trajectory analyses, comparisons of pseudotime-state occupancy between Low- and High-JDP2 groups were performed across the three biological replicates using paired statistical testing.

## Results

### Association between JDP2 expression and poor overall survival in colorectal cancer patients

To investigate whether metabolites, such as polyamines alter the expression of transcriptional regulators, we first performed a screening analysis using a panel of transcription factors. As part of the study, Caco-2 cells were treated with spermine for 24 h, followed by RNA isolation and qPCR analysis in a 96-well plate format (Supplemental Figure 1A; screening data not shown). Transcription factors showing altered expression in the initial screen were subsequently examined using different concentrations of spermine . Among the candidates tested, JDP2 showed a consistent reduction in expression following spermine treatment (Supplemental Figure 1B). JDP2 has been implicated in several malignancies, including T-cell acute lymphoblastic leukemia, liver hepatocellular carcinoma, kidney renal clear cell carcinoma, and uterine corpus endometrial carcinoma. Intriguingly, its association with tumor progression and patient outcome appears highly tissue-specific, tumor environment, and cell-type dependent^20–29^. For example, JDP2 has been shown to exhibit oncogenic and pro-survival functions in T-ALL but tumor-suppressive or favorable prognostic associations in several solid tumors. Despite this knowledge of JDP2 and cancer, the role of JDP2 in intestinal epithelial and colorectal cancer cell-cycle regulation remains understudied. To investigate the clinical implications of JDP2 in CRC, we first performed a Kaplan–Meier survival analysis (Figure 1) of the TCGA colon adenocarcinoma (TCGA-COAD) cohort. Patients were stratified by tumor JDP2 expression. As shown in Figure 1, we found that patients with higher JDP2 expression (dotted) have significantly worse survival than patients with lower JDP2 expression (solid) (log-rank P = 0.0112, Low JDP2, n = 227; High JDP2, n = 228). The survival curves showed progressive separation between the two groups, with the High-JDP2 group displaying a greater decline in survival probability over time. These findings demonstrate that elevated JDP2 expression is associated with an unfavorable clinical outcome in CRC and provide a rationale for investigating the functional role of JDP2 in colorectal epithelial cells.

**Figure 1.**
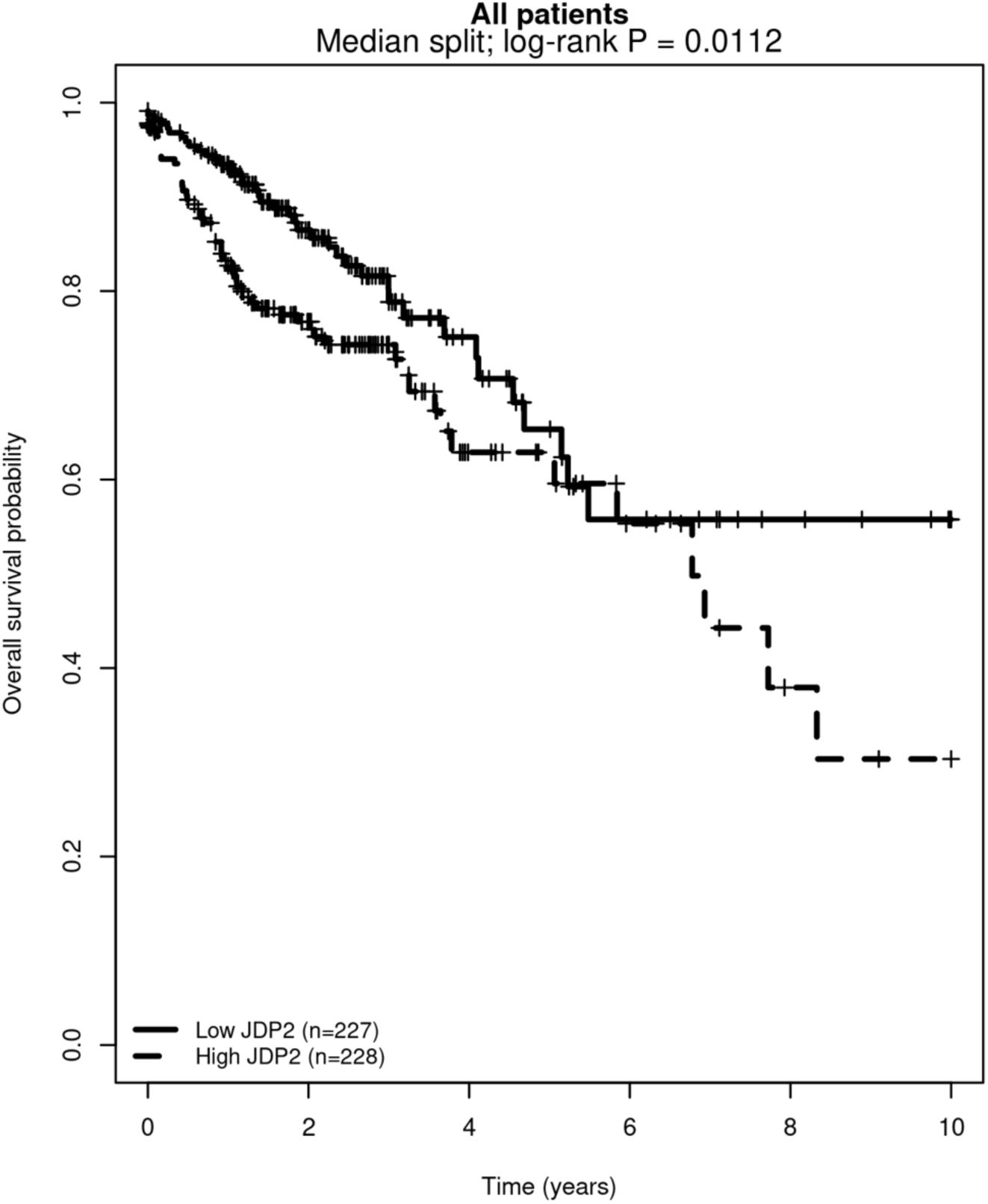
Association between JDP2 expression and poor overall survival in colorectal cancer patients. Kaplan–Meier survival analysis of TCGA-COAD CRC patients stratified by tumor JDP2 expression shows that patients with high JDP2 expression (dotted) have significantly worse survival than patients with low JDP2 expression (solid) (log-rank *P* = 0.0112, Low JDP2, n = 227; High JDP2, n = 228).

### JDP2 depletion alters cell-cycle progression in Caco-2 colorectal cancer cells

JDP2 has been reported to be associated with cancer in both oncogenic and tumor-suppressive roles, depending on the tumor type or cellular state^20–29^. To investigate the role of JDP2 in colorectal cancer cell proliferation, we performed siRNA-mediated JDP2 depletion in Caco-2 cells and examined cell cycle-associated markers. Based on the established functions of JDP2 in transcriptional and cell-cycle regulation, as well as its context-dependent roles in cancer type and cellular state, we hypothesized that depletion of JDP2 would alter cell-cycle progression in colorectal epithelial cancer cells.

For this purpose, we transfected Caco-2 cells at approximately 60-80% confluency with 12 pmol of control or JDP2-specific siRNA. At 72 hours post-transfection, we first assessed the efficiency of JDP2 knockdown by performing immunoblotting. Figure 2A shows that JDP2 protein abundance was markedly reduced (∼86% decrease in protein abundance) in cells transfected with JDP2 siRNA compared with cells treated with control siRNA. Furthermore, densitometric analysis also confirmed a significant reduction in JDP2 expression following siRNA treatment (Figure 2B).

**Figure 2.**
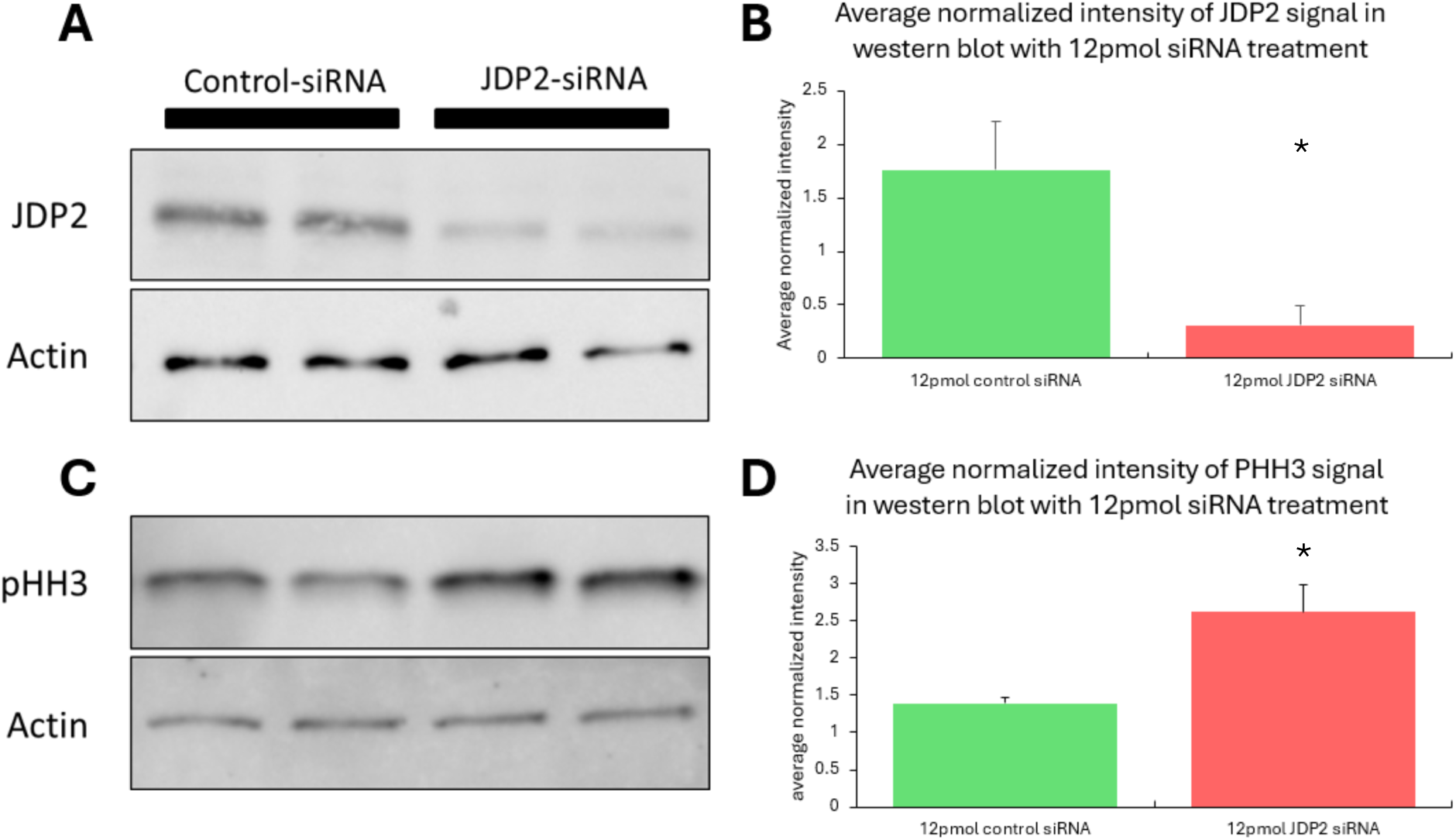
JDP2 knockdown increases phospho-histone H3 (PHH3) levels in Caco-2 cells. (**A**) Representative Western blot showing JDP2 protein levels in Caco-2 cells transfected with control siRNA or 12 pmol JDP2 siRNA. β-Actin was used as a loading control. (**B**) Average normalized intensity graph of JDP2 signal across multiple experiments after a 12pmol JDP2 siRNA treatment in Caco-2 cells. (**C**) Representative Western blot showing results of 12 pmol treatment of JDP2 siRNA on JDP2 level in Caco-2 cells. β-Actin was utilized as a loading control. (**D**) Average normalized intensity graphs of PHH3 signal across multiple experiments after a 12pmol JDP2 siRNA treatment in Caco-2 cells. PHH3 band intensity was normalized to β-actin, and data represent the mean normalized expression across independent experiments (P <0.05, t test).

Next, we examined phosphorylation of histone H-3 at serine-10 (PHH3), which is a marker associated with the cell cycle, specifically involved in chromosome condensation during late G2 and mitosis. We posited that depletion of JDP2 would alter the abundance of cells undergoing mitotic progression. As described above, at 72 hours post-transfection, we performed immunoblotting analysis shown in Figure 2C-D. JDP2-depleted cells demonstrated increased PHH3 signal compared with control siRNA treated cells (Figure 2C). Quantification across independent experiments confirmed a significant increase in actin normalized PHH3 signal following JDP2 depletion (Figure 2D). Overall, these findings suggest that JDP2 depletion by siRNA knockdown may alter colorectal cancer cell cycle progression and may be associated with an increased representation of cells exhibiting a late-G2/mitotic phenotype.

### JDP2 depletion increases the proportion of Caco-2 cells undergoing DNA synthesis

To further investigate the role of JDP2 in Caco-2 cell proliferation and cell-cycle activity, we evaluated DNA synthesis by EdU (5-ethynyl-2’-deoxyuridine) incorporation. EdU is a thymidine analog incorporated into newly synthesized DNA during S phase in place of endogenous thymidine and can be detected by a click-chemistry reaction. Thus, the EdU incorporation assay directly measures the fraction of cells undergoing active DNA synthesis during the labeling period.

In this assay, 72 hours post-transfection, Caco-2 cells were incubated with EdU for 2 hours, followed by fixation, EdU detection, and nuclear staining with DAPI. Figure 3A shows EdU-positive nuclei (red) and DAPI-positive nuclei (blue). The proportion of cells undergoing DNA synthesis was evaluated by calculating the ratio of EdU-positive cells to the total number of DAPI-positive nuclei.

**Figure 3.**
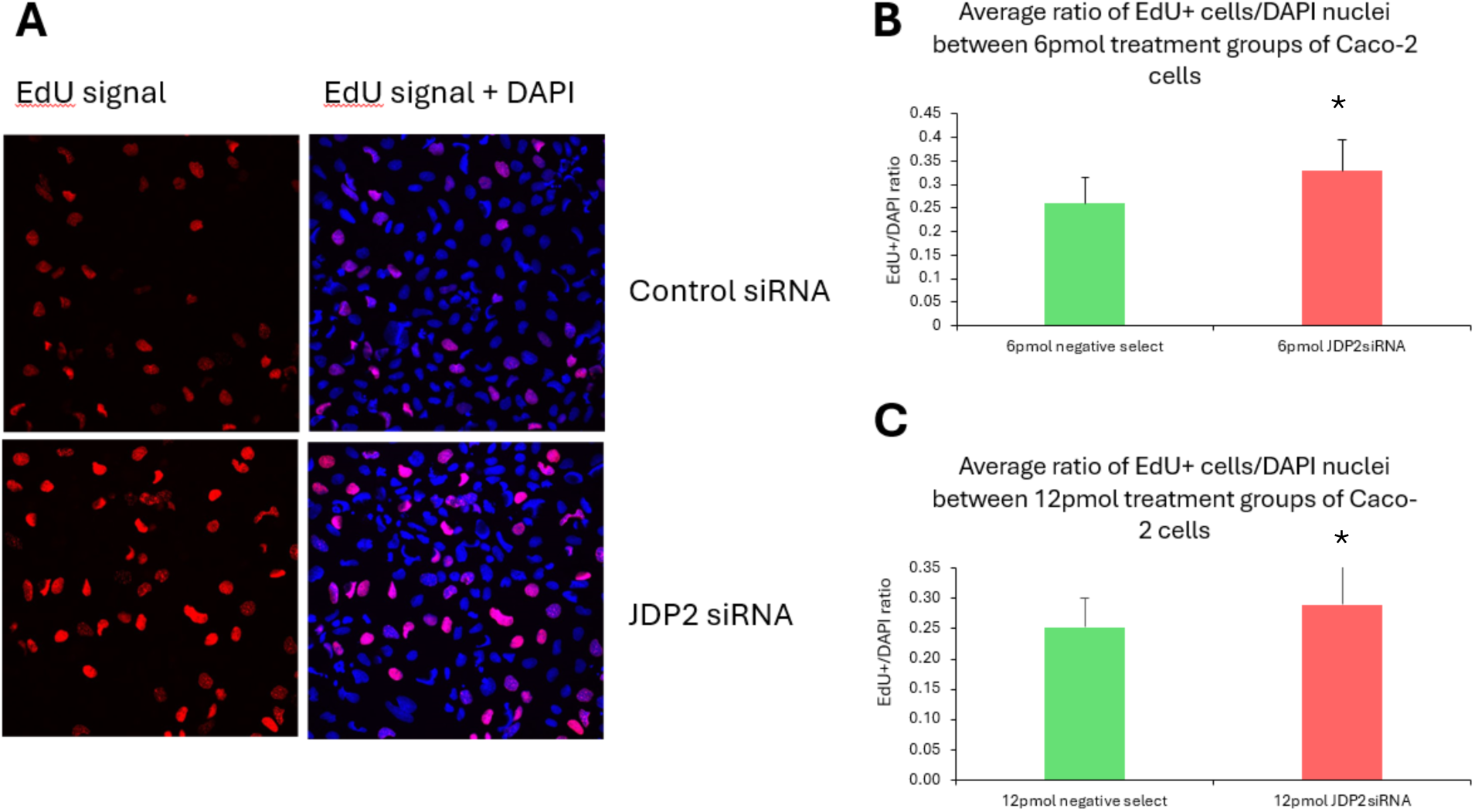
JDP2 modulates S-phase proliferation in Caco-2 cells. (A) Shown are representative fluorescence images of Caco-2 cells transfected with either control siRNA or 6 pmol JDP2 siRNA, then labeled for EdU incorporation. EdU-positive cells appear red, and nuclei are counterstained with DAPI (blue). The left panels display only the EdU signal, while the right panels show the merged EdU and DAPI signals. (B) Bar plot showing the average ratio of EdU-positive cells per DAPI-stained nucleus after treatment with 6 pmol JDP2 siRNA is shown. (C) Bar plot showing average ratio of EdU-positive cells relative to total DAPI-stained nuclei following treatment with control siRNA or 12 pmol JDP2 siRNA (P <0.05; t test).

At 12 pmol siRNA, JDP2 depletion significantly increased the Edu-positive/DAPI positive ratio compared to the control siRNA treated cells (Figure 3C), We also found that when Caco-2 cells were treated with 6 pmol of control or JDP2 siRNA, we still found a significant increase in the ratio of EdU incorporated cells in the JDP2 siRNA group (Figure 3B), which demonstrated that the effect was reproducible across two siRNA concentrations.

Taken together, the increased EdU-positive fraction and increased PHH3 abundance demonstrate that JDP2 depletion substantially alters cell-cycle dynamics in Caco-2 cells, which was characterized by increased abundance of cells undergoing both S-phase DNA synthesis and late G2/mitotic cell cycle progression. Hence, these findings suggest a role for endogenous JDP2 in restraining cell cycle progression in Caco-2 colorectal cancer cells.

### Single-cell RNA sequencing to reveal JDP2’s role in regulating IEC cell cycle homeostasis

To determine how JDP2 depletion affects cell-cycle-associated transcriptional states at single-cell resolution, we performed single-cell RNA sequencing using the 10x Genomics Chromium GEM-X Flex Fixed RNA Profiling platform with the Human Transcriptome Probe Set v2.0.0. Samples were processed as part of a multiplexed Flex experiment and demultiplexed using sample-specific probe barcodes. We processed sequencing data with Cell Ranger multi v10.0.0. We analyzed three independent JDP2-siRNA-treated Caco-2 samples (S1–S3; 91,187 cells). Our application of Unsupervised Seurat analysis identified 15 transcriptional clusters (UMAP plot, Figure 4A), and also determined cell-cycle scoring resolved G1-, S-, and G2/M-associated states across the transcriptional landscape (Figure 4B).

**Figure 4.**
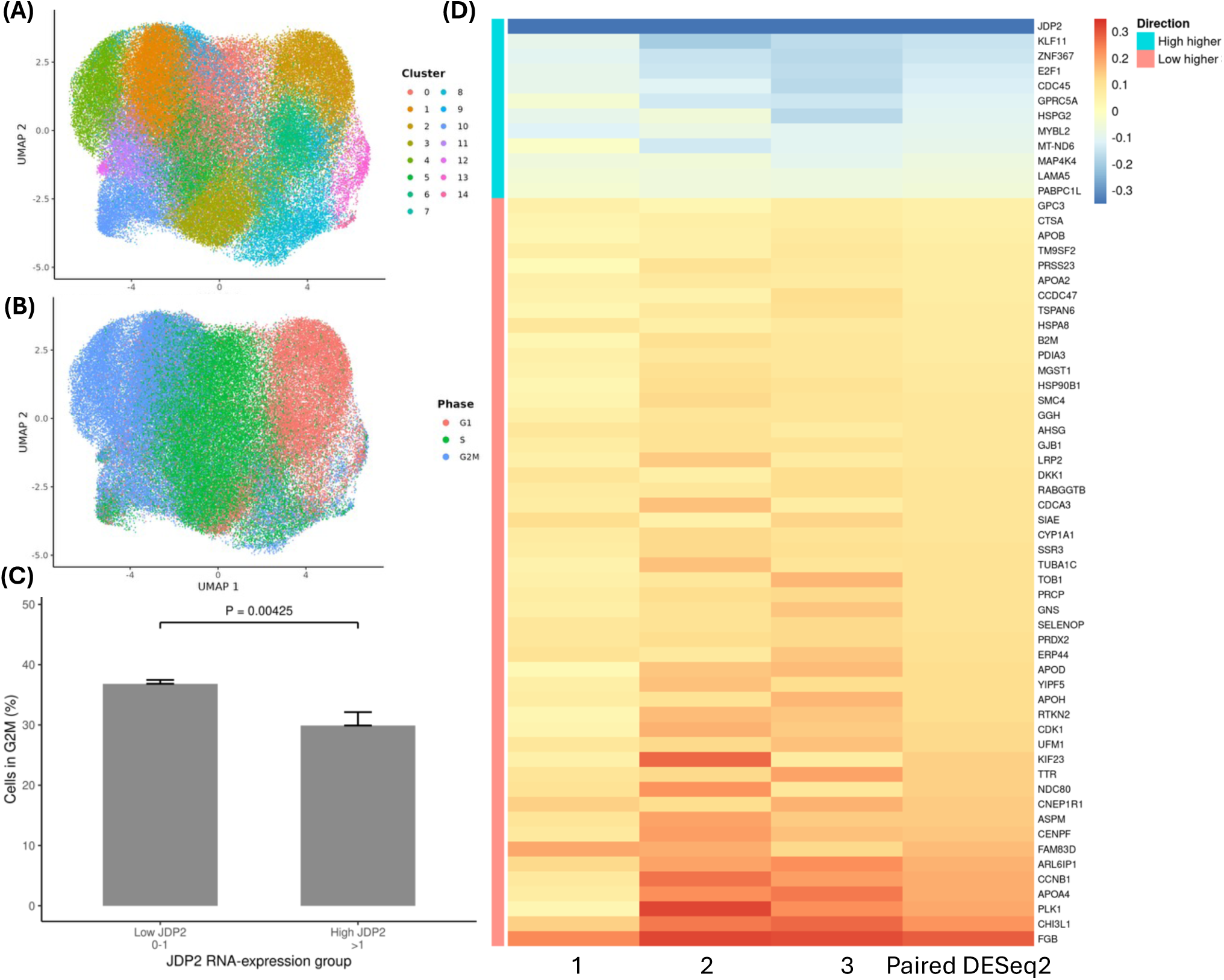
scRNA-seq analysis revealed that low residual JDP2 expression is associated with increased representation of the G2/M cell-cycle state in JDP2-siRNA-treated cells. We performed single-cell RNA-sequencing analysis on three JDP2-siRNA-treated biological replicates (n = 3). **(A)** A UMAP plot of the JDP2-siRNA-treated cells, with the cells colored according to the 15 Seurat transcriptional clusters. **(B)** A UMAP plot of the same cells, which had been assigned to the G1, S, and G2/M phases using Seurat cell-cycle scoring, illustrating the distribution of the various cell-cycle states throughout the transcriptional landscape. **(C)** A bar graph depicting the relationship between the amount of residual JDP2 RNA and the degree of G2/M representation, with the cell numbers balanced across the biological replicates; 2,000 cells were randomly selected from each replicate (giving a total of 6,000 cells) and divided according to their normalized JDP2 expression into Low (0–1) and High (>1) groups. The bars indicate the percentage of cells classified as G2/M. Cells with low JDP2 showed a higher degree of G2/M representation than those with High-JDP2 (36.83% compared with 29.91%). A binomial logistic regression found a significant association between the JDP2-expression group and G2/M status (comparing High to Low: OR = 0.732, 95% CI = 0.591–0.907, *P* = 0.00425). **(D)** A heatmap displaying the transcriptional differences that are associated with residual JDP2 expression among the JDP2-siRNA-treated cells (FDR <0.05).

JDP2 depletion by siRNA or genetic loss has been shown to alter cell-cycle progression and proliferation, including increased cyclin A2 expression and enhanced cell-cycle entry in mouse embryonic fibroblasts. JDP2 has also been examined using transcriptome-wide approaches after knockdown, demonstrating that JDP2 perturbation can produce cell state-dependent transcriptional responses. However, the role of JDP2 and its heterogeneity in expression in colonic epithelial cancer cells is not well studied. To model cell-to-cell heterogeneity in JDP2 expression analogous to that observed within tumors, we focused on the JDP2-siRNA-treated population and stratified individual cells according to residual JDP2 RNA abundance. As illustrated in Figure 2, immunoblot analysis revealed that JDP2-siRNA treatment led to a strong but incomplete JDP2 knockdown following JDP2-siRNA treatment (7.1-fold reduction; ∼86% decrease in protein abundance), leaving residual JDP2 expression within the treated population. Because heterogeneous gene expression is a common feature of tumor cell populations, we leveraged this residual cell-to-cell variation to model JDP2 expression heterogeneity. We therefore stratified JDP2-siRNA-treated cells according to residual JDP2 expression at single-cell resolution and examined whether JDP2-low and JDP2-high cells exhibited distinct transcriptional and cell-cycle states.

This design allowed us to determine whether JDP2-high and JDP2-low cells in the same cell-cycle compartment exhibited distinct transcriptional states, independent of initial siRNA exposure. To address heterogeneity in JDP2 expression among cells after siRNA treatment, we analyzed JDP2-siRNA-treated Caco-2 cells to determine whether residual JDP2 RNA expression was associated with cell-cycle state at single-cell resolution. We compared cells with Low (0–1) versus High (>1) normalized JDP2 expression using Seurat cell-cycle classification. This strategy allowed us to minimize unequal replicate representation. As shown in Figure 4C, Low-JDP2 cells demonstrated greater G2/M representation than High-JDP2 cells (36.83% vs. 29.91%), with a significant cell-level association (OR = 0.732, 95% CI 0.591–0.907, P = 0.00425). Furthermore, replicate-aware whole-population pseudobulk analysis identified a coherent mitotic transcriptional signature in Low-JDP2 cells, including increased expression of CDK1, CCNB1, PLK1, ASPM, CENPF, NDC80, CDCA3, SMC4, and KIF23 (Figure 4D). Thus, these findings suggest that lower residual JDP2 expression following JDP2 knockdown is associated with increased representation of the G2M/mitotic cell state (Figure 4).

As Figure 4 shows, our scRNA-seq analysis indicated that residual JDP2 expression defined High-JDP2 cells as a transcriptionally distinct state rather than a less proliferative counterpart of Low-JDP2 cells (Figure 4). Although Low-JDP2 cells showed greater G2/M representation and stronger late-mitotic transcriptional programs, High-JDP2 cells retained expression of cell-cycle and DNA-replication regulators, including E2F1, MYBL2, CDC45, and ZNF367, together with genes involved in signaling and extracellular-matrix interactions, including MAP4K4, HSPG2, and LAMA5 (Figure 4D). Among these gene expression signatures, LAMA5 is involved in basement-membrane organization and is associated with metastatic growth in CRC. MAP4K4 has been linked to tumor-cell migration, invasion, and other aggressive cancer phenotypes. HSPG2, a major component of the extracellular matrix, further points to differences in how High-JDP2 cells may interact with their surrounding environment. Taken together, these findings suggest that High-JDP2 cells are not simply a less proliferative version of Low-JDP2 cells but instead occupy a distinct transcriptional state with an ECM- and signaling-related component that may be relevant to tumor progression. Such non-proliferative properties could potentially contribute to the association between higher tumor JDP2 expression and poorer overall survival observed in CRC patients (Figure 1). Our research group is currently performing further studies to determine whether this state contributes to invasion or metastatic behavior and contributes to a metastatic or survival-altering phenotype.

### Pathway enrichment associated with residual JDP2 expression

We next investigated whether differences in residual JDP2 expression were associated with broader biological programs; we performed pathway enrichment analysis comparing Low- and High-JDP2 cells. Hallmark GSEA identified several pathways significantly enriched in Low-JDP2 cells (Figure 5A). These included the G2/M checkpoint and mitotic spindle, consistent with the greater representation of Low-JDP2 cells in late cell-cycle states. Interestingly, the differences included mTORC1 signaling, glycolysis, fatty-acid and bile-acid metabolism, peroxisomal activity, protein secretion, and xenobiotic metabolism. Hence, these findings indicate broader metabolic and signaling differences between the JDP2 expression states. We next restricted the analysis to G2/M cells to determine whether transcriptional differences remained when comparing cells within the same cell-cycle compartment (Figure 5B). Along this avenue, the Reactome analysis determined enrichment in Low-JDP2 cells, which included pathways related to IGF/IGFBP regulation, lipoprotein and LDL processing, integrin–MAPK signaling, sphingolipid and glycosphingolipid metabolism, and ER-to-Golgi protein transport. Thus, our study indicates that the differences between Low- and High-JDP2 cells were not limited to their relative distribution across cell-cycle phases, but were also associated with distinct metabolic and signaling programs.

**Figure 5.**
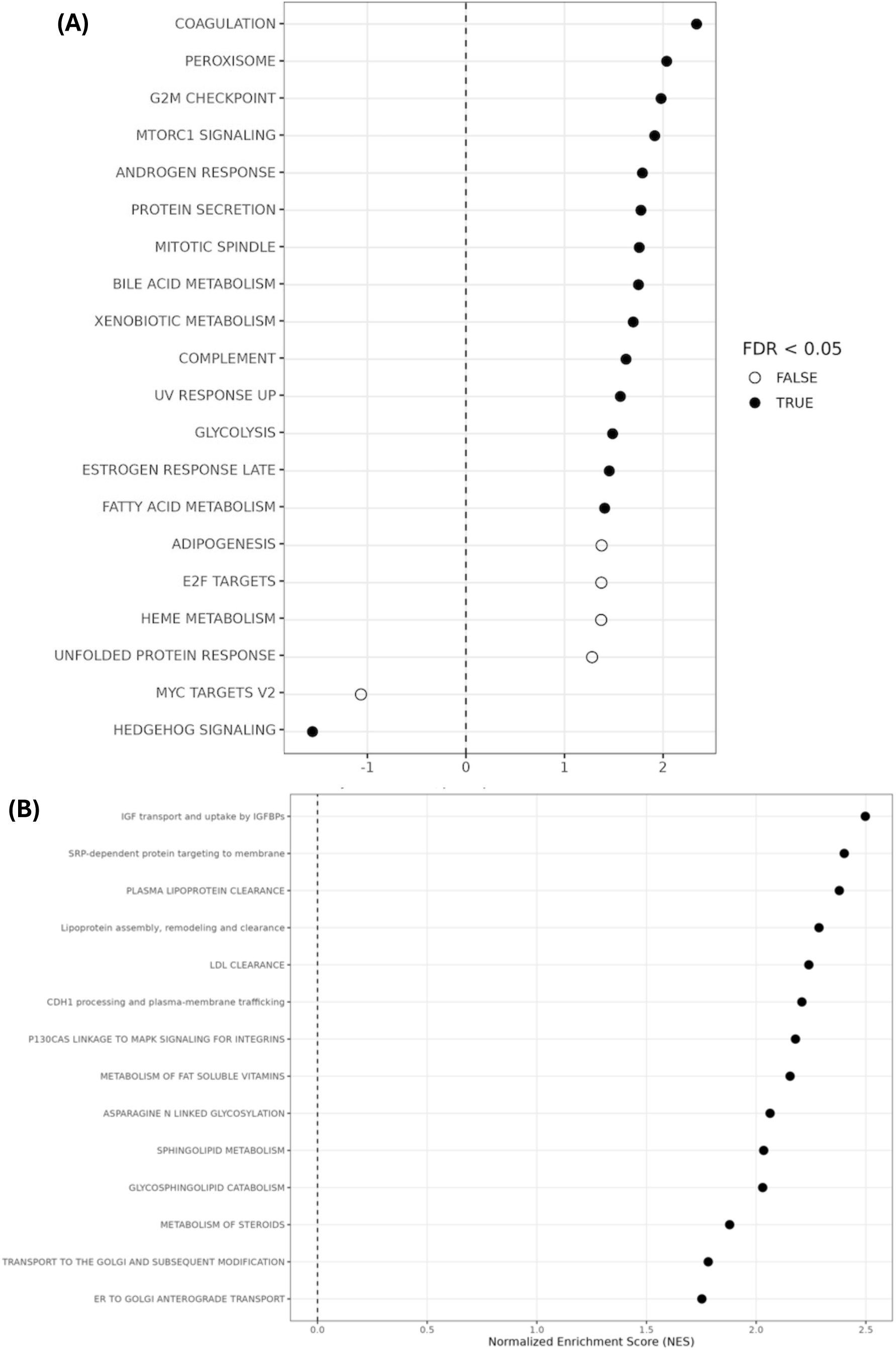
Pathway enrichment associated with residual JDP2 expression in JDP2-siRNA-treated cells. **(A)** Hallmark gene-set enrichment analysis comparing Low- and High-JDP2 cells across the three JDP2-siRNA biological replicates. Positive normalized enrichment scores (NES) indicate pathways enriched in Low-JDP2 cells, whereas negative NES values indicate enrichment in High-JDP2 cells. Filled circles indicate pathways meeting FDR < 0.05. **(B)** Reactome pathway analysis within the G2/M compartment using paired biological-replicate comparisons of Low-versus High-JDP2 cells revealed significantly enriched pathways in Low-JDP2 G2/M cells. Positive NES values indicate enrichment in Low-JDP2 cells.

### Residual JDP2 expression is associated with checkpoint and cell-fate heterogeneity within G2/M cells

Next, we wanted to investigate if the differences in transcriptional signature linked to JDP2 persisted even when we focused only on cells in the G2/M phase of the cell cycle (32,846 cells in total) (Figure 6). Within G2/M, canonical mitotic genes were not uniformly increased in Low-JDP2 cells. Consistent with this altered state occupancy, replicate-aware whole-population pseudobulk analysis identified a coherent mitotic transcriptional signature in Low-JDP2 cells, including increased expression of CDK1, CCNB1, and PLK1 (Figure 4D). In contrast, our analyses of G2/M-restricted violin and replicate-level expression revealed that the heterogeneity in checkpoint and cell-fate programs includes differences in CDC25C, CASP8, BIRC2, and GLB1 (Figure 6A). The replicate-by-group heatmap showed coordinated variation across checkpoint/mitotic, apoptotic, survival/adaptive, and stress/senescence-associated genes (Figure 6B). These findings suggest that the association between low residual JDP2 and G2/M representation is not explained by a uniform increase in canonical mitotic transcription and instead includes altered checkpoint and cell-fate states within the G2/M compartment.

**Figure 6.**
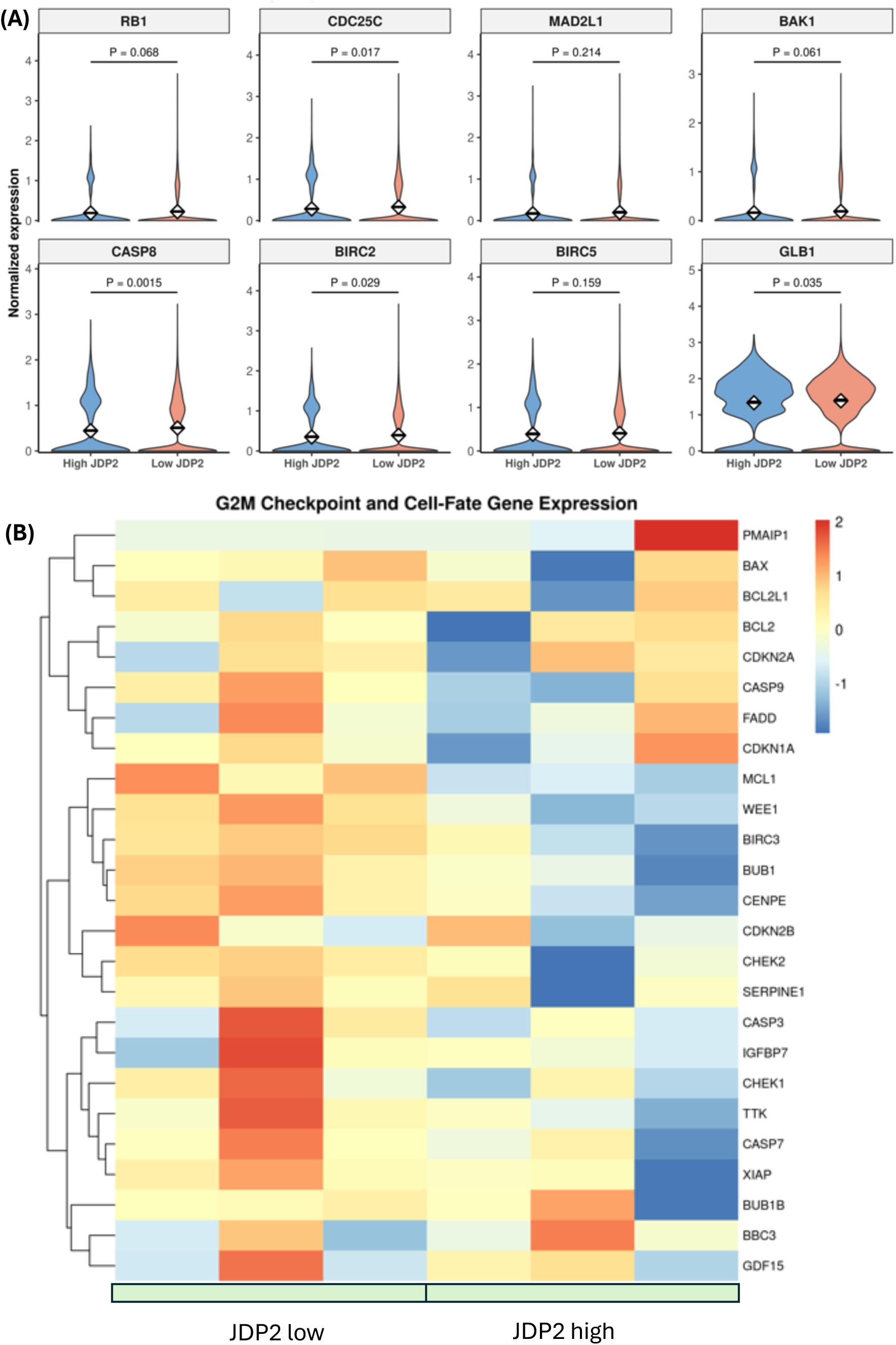
scRNA-seq analysis revealed that residual JDP2 expression is associated with transcriptional heterogeneity within G2/M-classified cells. To determine whether JDP2-associated transcriptional differences persisted within G2/M-classified cells, we restricted analyses to G2/M-classified cells from the three JDP2-siRNA biological replicates (n = 3). **(A)** Violin plots show normalized expression of representative G2/M checkpoint and cell-fate genes in High- and Low-JDP2 G2/M cells, including RB1, CDC25C, MAD2L1, BAK1, CASP8, BIRC2, BIRC5, and GLB1. **(B)** Heatmap of selected checkpoint/mitotic, cell-death, survival/adaptive, and stress/senescence-associated genes. We calculated mean normalized expression separately for each biological replicate and JDP2-expression group, then standardized each gene using row-wise z-score transformation; columns represent biological replicate/JDP2-expression combinations. We performed hierarchical clustering across genes, while keeping replicate/group columns unclustered.

### Low residual JDP2 expression is associated with increased occupancy of late cell-cycle-associated pseudotime states

To resolve cell-cycle-associated transcriptional progression beyond discrete phase labels, we applied Slingshot to the existing PCA representation of the JDP2-siRNA-treated cells. Our analysis revealed that Lineage 4 followed the cluster topology 2→6→3→5→12→4 and recapitulated an orderly G1→S→G2/M-associated continuum (Figure 7). The median Lineage-4 pseudotime increased from 26.22 in G1 cells to 36.47 in S cells and 44.40 in G2/M cells (Figure 7A). Pseudotime reflects a cell’s overall transcriptional state, not how much real time has passed. Interestingly, our analysis revealed that Low-JDP2 cells were shifted toward later Lineage-4 states in all three biological replicates. In addition, we subdivided pseudotime using Early, Middle, and Late boundaries. Intriguingly, our analysis determined that Late-state occupancy averaged 33.6% in Low-JDP2 cells, significantly more than 24.0% in High-JDP2 cells. This corresponded to a mean paired difference of +9.55 percentage points (paired t-test, P=0.048; Figure 7B). Furthermore, our replicate-resolved 20-bin occupancy profiles also showed the same redistribution toward later pseudotime regions (Figure 7C). Thus, our data revealed that the low residual JDP2 expression is associated with increased occupancy of late cell-cycle-associated pseudotime states.

**Figure 7.**
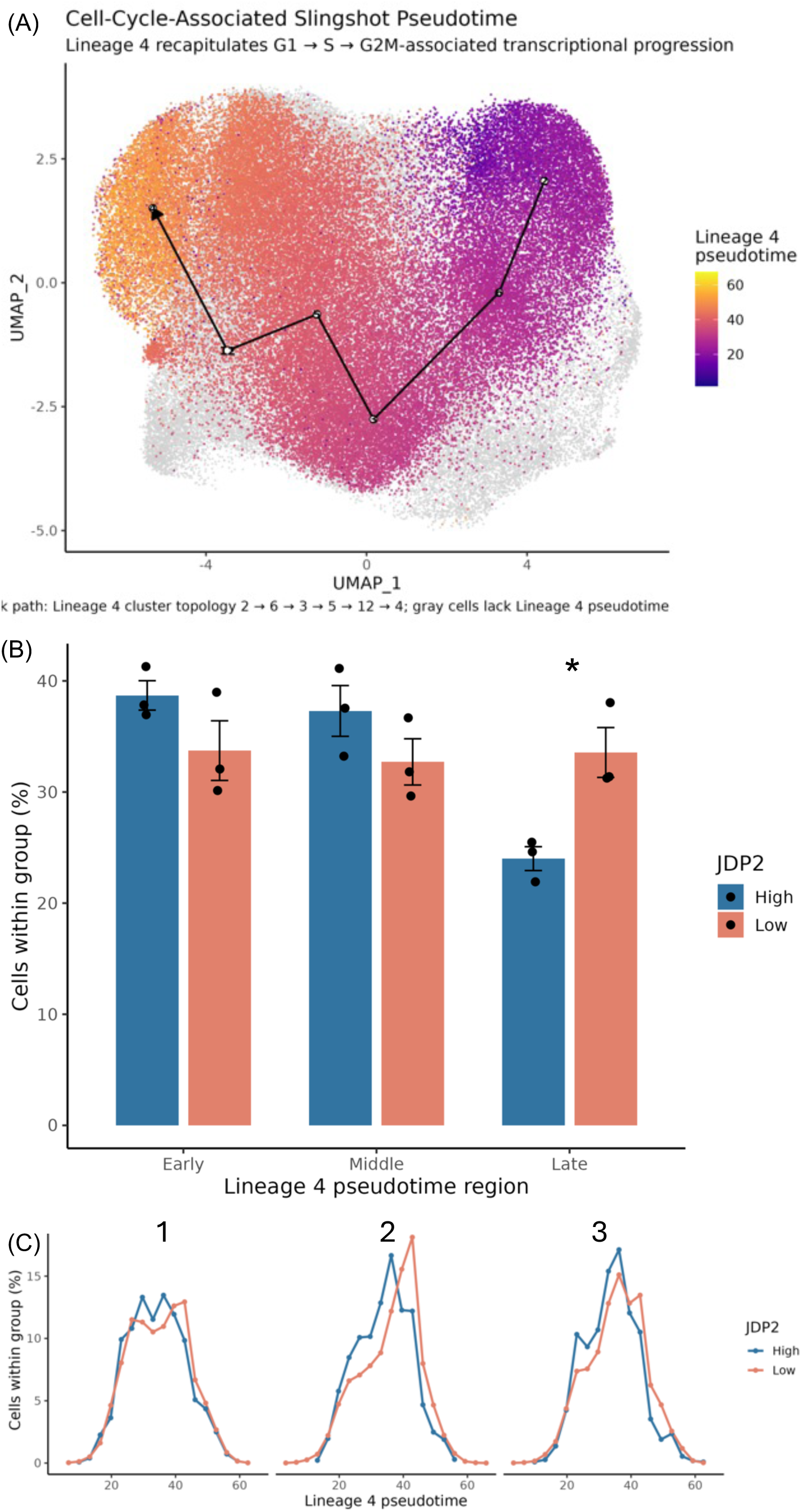
Low residual JDP2 expression is associated with increased late Lineage-4 occupancy. **(A)** Using Slingshot Lineage-4 pseudotime in JDP2-siRNA-treated Caco-2 cells, we recapitulated a G1→S→G2/M-associated transcriptional continuum (biological replicates, n = 3). **(B)** Bar plot showing Early, Middle, and Late pseudotime occupancy in Low-versus High-JDP2 cells. Low-JDP2 cells showed greater Late-state occupancy (33.6% vs. 24.0%; paired *P*=0.048). Bars represent mean ± SEM; points indicate biological replicates. **(C)** We analyzed normalized cell occupancy across 20 pseudotime bins for each biological replicate and found that Low-JDP2 cells were consistently more represented in later Lineage-4 pseudotime states. “Low JDP2” was operationally defined as log-normalized JDP2 expression ≤1, whereas “High JDP2” was defined as >1.

Taken together, our findings show that JDP2 depletion increases S-phase DNA synthesis and also late-G2/mitotic activity and shifts Caco-2 cells across distinct cell-cycle-associated transcriptional states. Single-cell analysis further revealed that cells with low residual JDP2 were more frequently represented in late pseudotime states and showed heterogeneity in checkpoint, survival, and cell-fate-associated programs rather than a uniform increase in mitotic gene expression. These findings suggest that JDP2 contributes to the regulation of colorectal epithelial cell-cycle homeostasis and that variation in JDP2 expression is associated with distinct proliferative and transcriptional states.

## Discussion

Our findings identify JDP2 as an important regulator of cell-cycle homeostasis in colorectal epithelial cells and reveal substantial heterogeneity in the transcriptional response to JDP2 depletion. Our data (Figure 2 - 3) demonstrated that siRNA-mediated JDP2 knockdown increased phosphorylation of histone H3 at Ser10 (PHH3) and EdU incorporation, indicating increased representation of cells undergoing late-G2/mitotic activity and S-phase DNA synthesis, respectively. These observations are consistent with previous reports that JDP2 can restrain cell-cycle progression^7–10^. Pan et al. reported that fibroblasts from JDP2-deficient mice showed increased proliferation and elevated cyclin A2 expression and demonstrated that JDP2 can repress the CCNA2 promoter^17^ (Pan et al., 2010). Cyclin A2 functions across S phase and G2/M progression. The report mentioned above supports a connection between JDP2 activity and the increased EdU incorporation and PHH3 signal observed in our experiments. Although our study does not establish cyclin A2 as the direct mediator of these effects in Caco-2 cells, the results are consistent with a model in which JDP2 contributes to the coordinated control of cell-cycle progression.

Our scRNA-seq transcriptomic analysis refines this interpretation by showing that JDP2 depletion is associated with redistribution across cell-cycle-associated transcriptional states rather than a simple uniform increase in a mitotic program. We examined 91,187 JDP2-siRNA-treated cells; and we found that Low-JDP2 cells showed greater representation of the G2/M state than High-JDP2 cells. Consistent with this altered state occupancy, replicate-aware whole-population pseudobulk analysis identified a coherent mitotic transcriptional signature in Low-JDP2 cells, including increased expression of CDK1, CCNB1, PLK1, ASPM, CENPF, NDC80, CDCA3, SMC4, and KIF23. Furthermore, Hallmark GSEA identified several pathways significantly enriched in Low-JDP2 cells (Figure 5A), including the G2/M checkpoint and the mitotic spindle as signature pathway enrichments.

Our scRNA-seq analysis data further demonstrated that Low- and High-JDP2 cells within G2/M exhibited heterogeneity in checkpoint, apoptotic, survival, and stress-associated programs. Notably, in the late-Lineage-4 analysis (Figure 7), we found an increase in several checkpoint-associated transcripts in Low-JDP2 cells, including MDM2, WEE1, and ATR at nominal paired significance. Furthermore, our analysis identified BIRC2 as a consistent Low-JDP2-associated increase in the initial late-state gene panel. Together, these results favor a model in which reduced residual JDP2 is associated with altered checkpoint and cell-fate regulation within late cell-cycle states rather than simply stronger canonical mitotic transcription.

Across cancer models, the impacts of JDP2 appear to be variable among tumor types and cell-state-specific^20–29^. In our findings in colorectal cancer cells, JDP2 functions as a negative regulator of proliferation. This aligns with findings from Heinrich et al., who demonstrated that JDP2 was tumor suppressive and inhibited cell transformation in a prostate cancer cell line^11^ (Heinrich et al., 2003). In contrast, Bitton-Worms et al. demonstrated that JDP2 transgenic mice displayed potentiation of liver cancer and that JDP2 expression at the promotion stage was most critical for increasing severity of liver cancer^12^ (Bitton-Worms et al., 2010). The difference in findings between the role of JDP2 in liver cancer, leukemia, and prostate cancer highlights the need for further study on the role of JDP2 in various cancers, including colorectal cancer^20–29^.

To investigate the role of JDP2 in the colorectal cell cycle, we have performed scRNA sequencing analysis instead of bulk RNA analysis. Our JDP2 siRNA treatment of Caco-2 cells and Western blot analysis confirmed residual JDP2 even after siRNA treatment (Figure 2). To evaluate the effect of residual JDP2, we performed scRNA-seq gene expression analysis across cell populations, because bulk RNA sequencing analysis can obscure heterogeneous expression of a target gene associated with distinct transcriptional states within the same cell population. Bulk RNA-seq is instrumental for studying population-level gene expression; however, it masks valuable individual cell transcriptome status in disease conditions or tissue microenvironments.

This distinction is especially relevant to cancer. Tumors contain malignant cells occupying multiple transcriptional states even within the same tumor and therefore sharing the same patient-level clinical context. Patel et al. demonstrated extensive cell-to-cell transcriptional heterogeneity within individual glioblastomas, including variation in proliferation and other oncogenic programs^31–36^. Tirosh et al. similarly showed that malignant cells from the same melanoma tumor differed in cell-cycle state and drug-resistance programs. Interestingly, tumors characterized by high MITF still contained individual MITF-low/AXL-high malignant cells. Thus, a healthy control tissue is not required to ask an intratumoral heterogeneity question: the biological comparison can be between malignant-cell states existing within the same tumor. These studies illustrate a major advantage of single-cell analysis: individual malignant cells within that tumor can be studied and positioned along transcriptional states or cell cycle programs without requiring each state to have a matched normal counterpart.

Trajectory analysis gave another perspective on this redistribution of states. Slingshot Lineage 4 recapitulated a G1→S→G2/M-associated transcriptional continuum, and Low-JDP2 cells showed increased occupancy of its late region in all three biological replicates (Figure 7). The level of late-state occupancy was about 33.6% in the Low-JDP2 cells as against 24.0% in the High-JDP2 cells. Our result should be interpreted as differential occupancy of transcriptional states. Therefore, it should not be interpreted as direct evidence of altered cell-cycle velocity, accumulation, or arrest, because pseudotime orders cells by transcriptional similarity and does not measure elapsed biological time. Nonetheless, the agreement between EdU incorporation, PHH3 abundance, along with discrete cell-cycle classification, and pseudotime occupancy supports our broader conclusion that JDP2 depletion perturbs the balance of proliferative cell states.

The biological functions of JDP2 vary considerably across cell types and biological settings. JDP2 has been reported to suppress transformation and cell-cycle progression in some systems, whereas pro-survival or tumor-promoting functions have been described in others, including T-cell acute lymphoblastic leukemia and hepatocellular carcinoma^11–29^. This context dependence is also relevant to our TCGA-COAD analysis, in which higher tumor JDP2 expression was associated with worse overall survival (Figure 1). The clinical association does not contradict the increased proliferative activity observed after acute JDP2 depletion in Caco-2 cells: tumor JDP2 abundance may reflect additional functions in survival, stress adaptation, differentiation, or tumor-state composition that are not reproduced by acute siRNA depletion in a cultured epithelial model. Furthermore, in our scRNA-seq analysis, we found that the High-JDP2 cells represented a transcriptionally distinct state rather than simply a less proliferative counterpart of Low-JDP2 cells. Although Low-JDP2 cells showed greater G2/M representation and stronger late-mitotic transcriptional programs, High-JDP2 cells retained expression of cell-cycle and DNA-replication regulators, including E2F1, MYBL2, CDC45, and ZNF367, together with genes involved in signaling and extracellular-matrix interactions, including MAP4K4, HSPG2, and LAMA5 (Figure 4). Interestingly, of the High-JDP2 signature, LAMA5 and MAP4K4 are particularly relevant to tumor progression. LAMA5 relates to basement-membrane/ECM biology and has been implicated in CRC metastatic growth. Additionally, MAP4K4 has been linked to migration/invasion and aggressive cancer phenotypes. HSPG2, on the other hand, also points toward altered tumor– ECM interactions. Thus, High-JDP2 cell contains a biologically coherent ECM/signaling/tumor-progression component rather than being simply a low-proliferation population. Therefore, within the G2/M compartment, differences in checkpoint, apoptotic, and cell-fate-associated genes further indicated that JDP2-associated transcriptional heterogeneity persisted independently of differences in cell-cycle-phase occupancy. These findings highlighted that High-JDP2 cells occupy a qualitatively different cellular state rather than simply displaying an inverse mitotic phenotype. Interestingly, the signaling and extracellular-matrix components of the High-JDP2 signature raise the possibility that higher JDP2 expression may accompany tumor-cell properties related to interactions with the tumor microenvironment, invasion, or metastatic progression rather than simply increasing proliferation. However, further studies are needed to determine whether these transcriptional differences translate into functional changes in tumor-cell survival, invasion, or metastatic potential. Examining JDP2-associated states in patient-derived tumor tissues, together with functional and in vivo studies, will be particularly important for determining whether these properties help explain the association between higher tumor JDP2 expression and poorer patient survival.

In summary, our data demonstrated that JDP2 depletion increased S-phase DNA synthesis and late-G2/mitotic markers and redistributed Caco-2 cells across cell-cycle-associated transcriptional states. Interestingly, single-cell RNA sequencing analysis further revealed that low residual JDP2 expression was associated with increased occupancy of late pseudotime states and with checkpoint and cell-fate heterogeneity that was not explained by a uniform increase in canonical mitotic gene expression. Hence, our study supports a role for JDP2 in maintaining colorectal epithelial cell-cycle homeostasis and establishes a single-cell perturbation framework for investigating target-gene-associated transcriptional heterogeneity in colorectal cancer.

## Author Contributions

M.A.A. conceived and designed the study, supervised the research, acquired funding, and contributed to data interpretation and manuscript preparation. V.N., A.G., A.E., M.F., and P.B. contributed to experimental investigation, data collection, and analysis. M.A.A. performed the single-cell RNA-sequencing bioinformatic and computational analyses and integrated the experimental and single-cell findings. V.N., A.G., and M.A.A. prepared the initial manuscript draft.

---

**Supplemental Figure 1.**
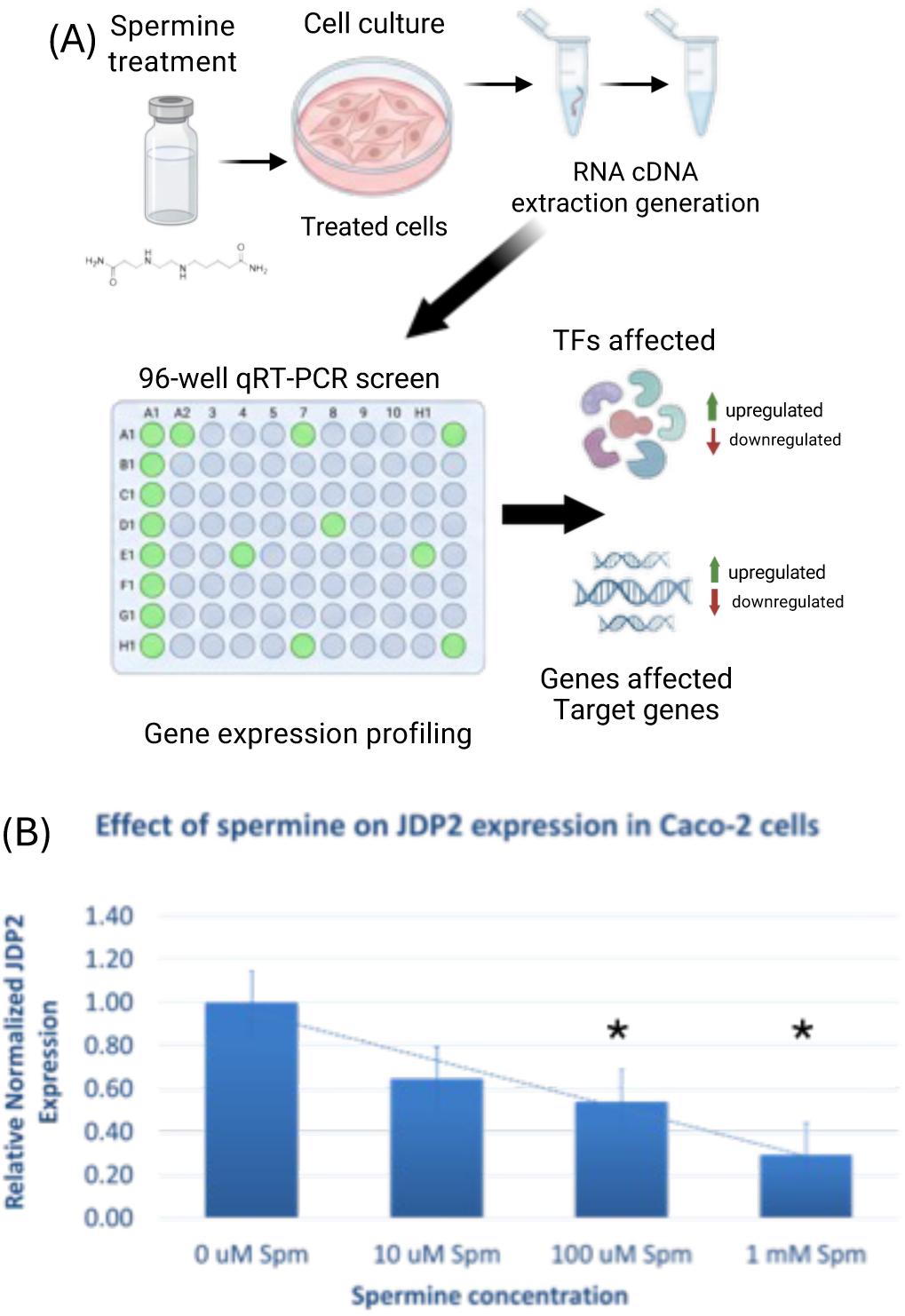
The effect of spermine on relative JDP2 expression in Caco-2 cells. (A) Schematic overview of the experimental approach used to identify transcription factors whose expression is altered by polyamines. (B) Caco-2 cells were incubated with increasing concentrations of spermine (0 *μ* M to 1000 *μ*M). Extracted RNA was used to perform RT-qPCR using the iTaq Universal SYBR Green One-Step Kit. Data are presented as relative normalized JDP2 expression (mean ± SEM, *P < 0.05).

